# Automated morphometry and weight prediction of juvenile Chinook Salmon leveraging open-source deep learning models

**DOI:** 10.64898/2026.03.10.710725

**Authors:** Brian Knight, Carson Jeffres

## Abstract

Minimizing handling of threatened and endangered fish has become increasingly important as populations have dwindled. To minimize handling in morphometric measurements, the HandsFreeFishing program has been developed for juvenile Chinook Salmon (Oncorhynchus tshawytscha). By segmenting a 2D image many morphometric measurements are able to be estimated; from these measurements a weight prediction model is built based on fish whose ground truthed weights were measured using a digital scale. While many segmentation methods may be used, here Meta’s Segment Anything model (SAM) is employed to produce segmentation masks of raw images. This model is open-source and easily used on any image (of any size) with good performance. In the proposed framework, the user supplies a bounding box around a target fish along with minimal orientation data (left or right facing, upside down or right-side up); the rest of the segmentation, feature extraction, and final weight prediction is completely automated. A main goal of the segmentation is to estimate the surface area of the side profile of the fish. Then, assuming an ellipsoidal shape, this surface area can be related to the volume of the fish, which is directly proportional to the weight. Even on a relatively small dataset of 149 images (fork length 27-90mm) our results confirm the predictive qualities of the morphometric features measured. The model achieved weight prediction with a mean absolute error of 0.16 g with a mean absolute percentage error of 12%, and an *r*-squared value of 0.99, on fish ranging from 0.31g - 7.74g. The raw images come from a variety of fish viewers, the design of which is relatively inexpensive and reproducible, and, in conjunction with the HandsFreeFishing program, allows for minimal handling compared to traditional length and weight measurement methods.

## Introduction

Chinook Salmon (*Oncorhynchus tshawytscha*) populations along the California coast are struggling due to many factors including overfishing and loss of habitat. These fish are an important source of food both commercially and for recreational fisheries. Further, Chinook Salmon are important for Indigineous communities, historically serving as an vital source of food and cultural signifcance. Juvenile Chinook Salmon rearing and migratory habitat has been degraded, limiting the opportunity for the expression of diverse of life history strategies. (Kareiva et al. 2000 [1], David et al. 2016 [2]). There is December 3, 2025 1/10 a large ongoing effort to restore these habitats and fish populations to healthy levels including dam removal projects, habitat restoration, and novel management actions. Further, it is known that certain morphometric measurements of juvenile fish are indicative of overall fish health, such as Fulton’s condition factor (*K*) defined as *K* = 10^5^ × *W* × *L*^−3^, where W is weight and L is length (Ricker 1975 [3]). The work described here is important for this effort in two majors ways. First, it proposes an practical platform to measure accurate morphometric data of individual fish based on a side-profile image, allowing for fast and consistent measurement in many environments. Second, it is designed specifically to minimize fish handling, keeping the fish submerged in water throughout, directly addresses a growing need for sustainable and non-invasive methods to protect fish health during measurement (Sharpe et al. 1998 [4]).

In 2021 Holmes and Jeffres [5] proposed a morphometric model for weight prediction and condition factor prediction for juvenile Chinook Salmon as manual weighing and measuring of individual juveniles is challenging and time consuming work, and is prone to user error. However, the time required to digitize each morphometric point on every fish remains a bottleneck in the data processing pipeline, and, moreover, must be done by a trained user. Here, to collect more accurate and reproducible information, as well as minimize handling of the fish, the HandsFreeFishing program has been developed, which uses an open-source deep-learning based segmentation models to automate much of the data processing pipeline. This program requires minimal user input for each individual image, while allowing researchers to measure important morphometric quantities of interest. Meta’s open-source Segment Anything Model (SAM) [6] was used here in order to segment individual fish, then customizable algorithms were applied to extract relevant morphometric features, such as fork length and surface area, which were later eventually used to predict fish weight. There have been several previous attempts to predict fish weight using image data and deep learning. Yang et al. [7] (2021) built a deep convolutional neural network to predict the weight of fish through 2D image data. Rantung, Sappu and Tondok [8] (2021) used stereo vision, requiring several images of a fish from different angles, to predict fish surface area and volume. The approach developed here is more similar to the [8], as we focus on estimating well defined morphometric features which are then used to predict fish weight, though we use only one side-profile image per fish, as done in [7]. Lastly, another automatic morphometric model was proposed by Kristiansen et al. in 2025 [9], which trains a machine learning model to place landmark points of zebrafish. In this work, training data is used only to predict fish weight; all other morphometric features are computed directly from the fish segmentation and contour.

All of this study was focused on our juvenile Chinook Salmon (fork length 27-90mm and weight 0.31g - 7.74g), but these programs are open-source and may be easily customized to extract morphometric features from image data of other species as well.

## Materials and methods

A conceptual overview of the methodology is presented in Fig 1. All necessary software to reproduce these results can be found at https://github.com/briancknight/HandsFreeFishing, where installation steps and examples are outlined. It requires Python 3.10 or later, and access to Meta’s open-source Segment Anything Model (SAM) [6].

**Fig 1.**
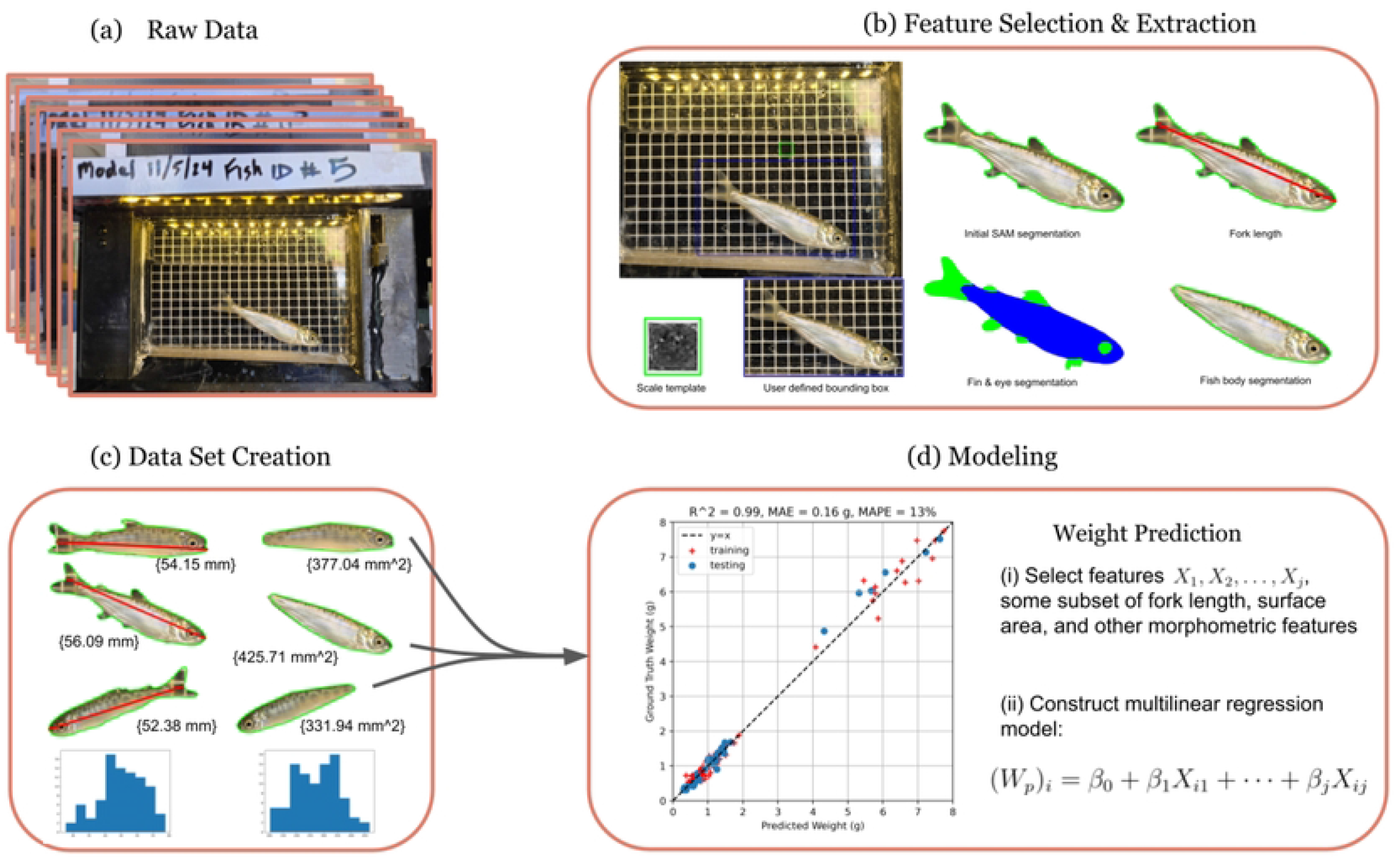
A conceptual overview of the proposed data processing pipeline

### 0.1 Image Collection

All fish images were acquired using a point and shoot digital camera of fish inside a viewer developed to collect images of live juvenile salmon up to 90mm (Fig 1). The viewer held fish in water at a side profile where the back side of the viewer contained a 5×5mm grid used for standardizing a digital length in the images. The viewer contained lights above the viewing chamber to illuminate the fish to ensure fast enough shutter speeds to collect sharp, in focus images. Following image acquisition, the fish were returned to a holding bucket and ultimately returned to their rearing tank.

### 0.2 Initial Image Processing

Once aggregated, each image was manually inspected and given a digital bounding box and orientation (left facing or right facing, upside down or right-side up). The overall image quality was assessed at this stage too, to ensure any unusable images were discarded. An example of this interface is shown in Fig 2 (a)(b). The user interface is designed to be simple, reading each image name in from a corresponding spreadsheet or csv file.

**Fig 2.** (a) Raw image displayed by our program, waiting for a user-supplied bounding box. (b) A bounding box is supplied by the user with two clicks. Once the bounding box is confirmed, the user is asked for minimal orientation information regarding the fish. (c) The raw image (d) The initial segmentation via SAM.

### 0.3 Automated Segmentation

After initial image processing, Meta’s Segment Anything Mode (SAM) [6] is used to extract an initial segmentation of the target fish, as shown in Fig 2(c)(d). Further, a smooth contour of the fish outline is computed using elliptic Fourier analysis, which yields a discretized smooth 2D curve *c* = [*c*_0_, *c*_1_, …, *c*_*n*_] where *c*_*i*_ = (*x*_*i*_, *y*_*i*_). The mesh grid from the fish viewer (in the background of each image) is used to estimate the pixel to millimeter scale. This estimate is done by template matching, finding a grid box which we know to represent a 5 by 5-millimeter square. Using the initial segmentation, contour and scale estimate, the program estimated the size location of individual fins and again applied SAM with bounding boxes computed from these estimates. Examples of successful and unsuccessful segmentations from this procedure are shown in Fig 4(a)(b). With individual fins segmented, they are then removed from the initial segmentation, yielding a ‘no fin’ segmentation (Fig 4(d)). Because fish fins can be in several different positions (up or down or in-between), they can influence the side-profile surface area which can affect weight prediction. As such it has been determined to remove fin segmentations [5]. In addition, the eye of the fish was segmented to measure eye diameter, a potentially important morphometric measurement.

### 0.4 Feature Extraction

For all feature extraction, the fish is assumed to be right-side up and facing left. The orientation data supplied by the user ensures each image is rotated and flipped appropriately, so that the program can process each contour in the same fashion.

#### 0.4.1 Fork Length & Surface Area Measurement

The fork length, the distance from the tip of the jaw or snout with closed mouth to the center of the fork in the tail, is a standard measurement in fish ecology. To measure the fork length of each fish, the shape contour is first rotated so that the left-most point and right-most point lay along a horizontal line. After rotation, the new left-most point of the contour is taken to be the tip of the fish’s head. To find the correct point of the caudal fin the program searched along the opposite end of the contour for two consecutive local maxima in the *x*-component which would represent the upper and lower lobes of the caudal fin; these contour point are called 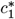 and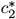. If these points are found successfully (Fig 3(a)-(c)), then the program now has a segment of the contour 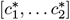 within which it will find the point with minimum *x*-component. If it cannot find the upper and lower lobe points (Fig 3(d)), it assumes the caudal fin is not appropriately segmented, and instead does the following: consider only the contour points whose *x*-component is larger than some threshold (this gives only the contour of tail-end of the fish). Then, compute the average *y*-component (height) of these contour points, and find the point *c*^∗^ whose height is closest to this average height. This should be near where the midpoint of the caudal fin would be given a proper segmentation.

**Fig 3.** Fork length predictions from four separate images following the procedure outline in section 0.4.1.

The automated program uses similar style procedures for segmenting individual fins: caudal, adipose, dorsal, pectoral, pelvic, and anal, as well as for segmenting the eye of the fish. For high quality images, this produces good fin segmentations which can be used to measure various morphometrics researchers may be interested in. Details on the heuristics we use for each of these are included in the S1 Appendix 0.5, but these heuristics are customizable, which is a feature of the program. Fig 4 shows some examples of successful and unsuccessful fin segmentation, and corresponding ‘no fin’ fin segmentations. The ‘no fin’ segmentations are of interest because are highly correlated with the fish volume, which is directly related to fish weight via density. For more discussion on weight prediction, see Section 0.5. If fin segmentation is reliable, the individual surface area measurements could be used to improve a weight prediction model, or could be measured for other purposes. In this case the fin segments were removed from segmentation, as the fish can move them up or down resulting in less reliable segmentation. Additionally, because the fin density varies drastically from the density of the rest of the body it can lead to erroneous weight predictions. To remove the fins from the segmentation, masked areas were set equal to zero. When this step is performed, the resulting segmentation may have multiple connected components, or, when fin segmentation is unsuccessful, may remove too much area from the body of the fish. To mitigate these issues, two steps are performed. First, only the largest connected component of the segmentation is kept. Second, using the contour of the segmentation, a convex hull of the contour points is computed. So long as the area of the convex hull is not significantly larger than the area of the segmentation, this convex hull is kept as the final segmentation used in further processing.

**Fig 4.**
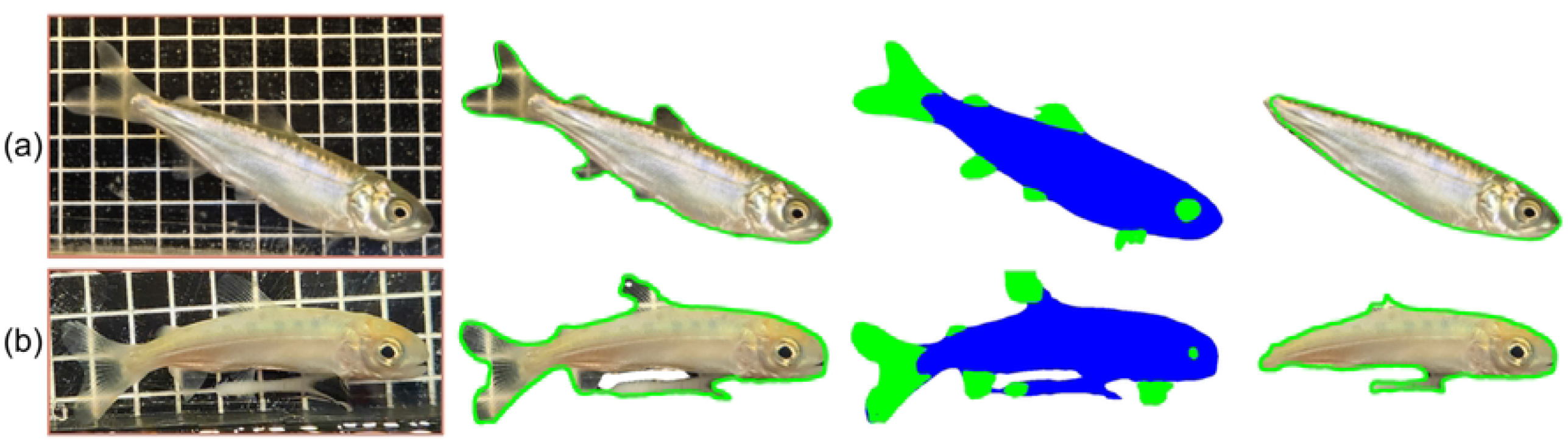
Two examples of automated segmentation and contour estimation after initial bounding box: (a) successful, (b) unsuccessful.

For the remainder of the paper, fish segmentation and fish contour will refer to the ‘no fin’ segmentation and ‘no fin’ contour respectively (e.g. the right most image of Fig 4(a)).

#### 0.4.2 Other morphometric features

In addition to the fork length and surface area estimates, the program also computed eye diameter based on the eye segmentation mentioned previously, as shown in Fig 5(b). The estimated diameter is computed as the maximum distance between any two points along the contour of the eye segmentation.

**Fig 5.**
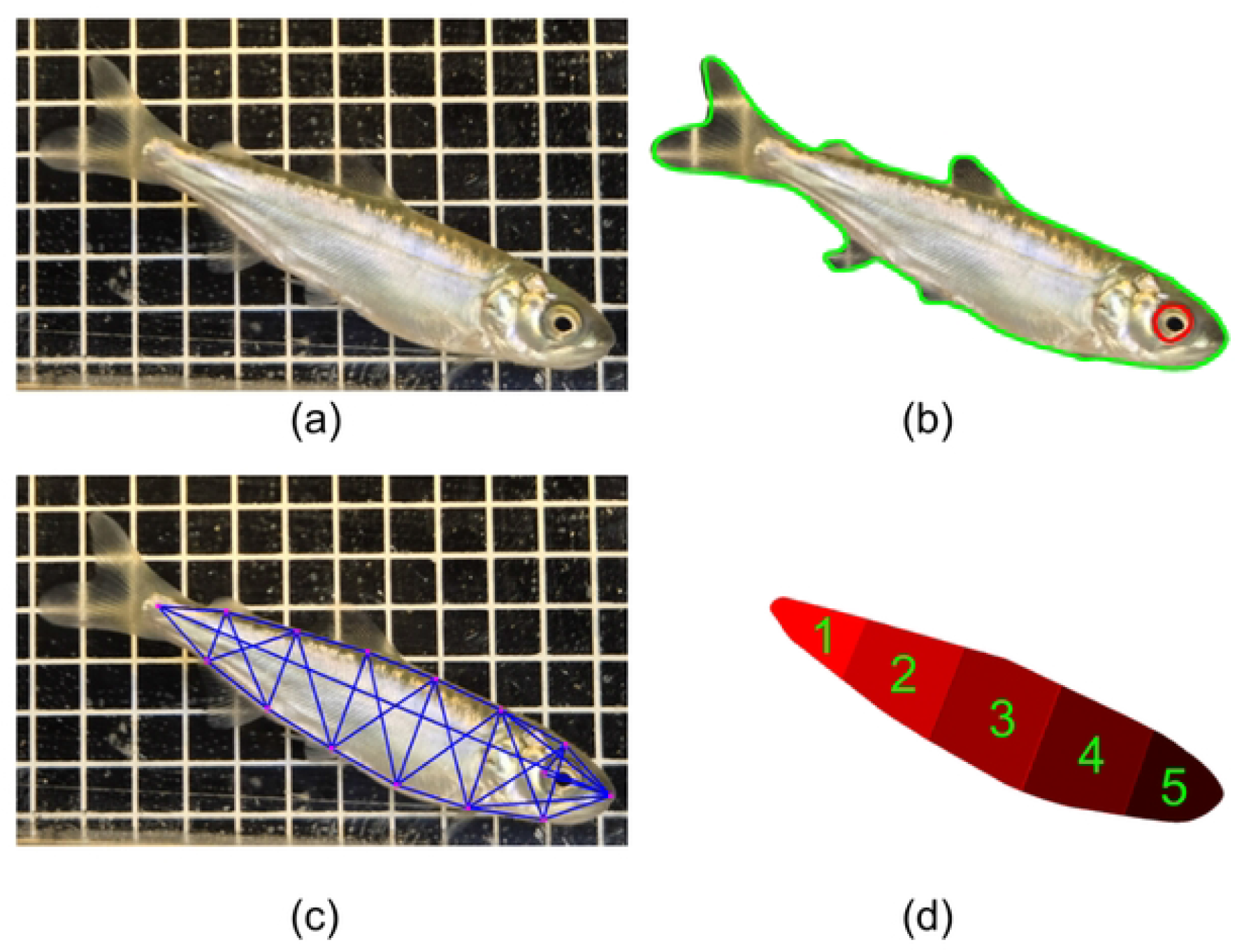
Other morphometric features: (a) raw image, cropped, (b) eye contour used for eye diameter, (c) landmark points & lengths (d) partitioned surface area.

To compare previously described morphometric modeling [5], the program automated the placement 16 morphometric points the fish image. In contrast to previous work (Holmes and Jeffres 2021), these landmark points were not based upon fin locations, but instead placed at 6 equally spaced points in between the left and right most point of the contour, along with a point just behind the eye. A truss lattice was then constructed between these points, and the length of each of the line segments shown in Fig 5(c) were measured. Even without the correct anatomical placement, these length measurements provide similar data about the size and shape of the fish.

Additionally, the program can compute partitioned surface areas, which could allow a model to learn a non-constant density function across the body of the fish (Fig 5(d)). Lastly, we found the ellipse of best fit through the points defining our contour, and measured the major and minor semi-axes, *a* and *b*, of this ellipse, which are proximal measurements for the length and height of the fish respectively. A visualization of these quantities is shown in Fig 6.

**Fig 6.**
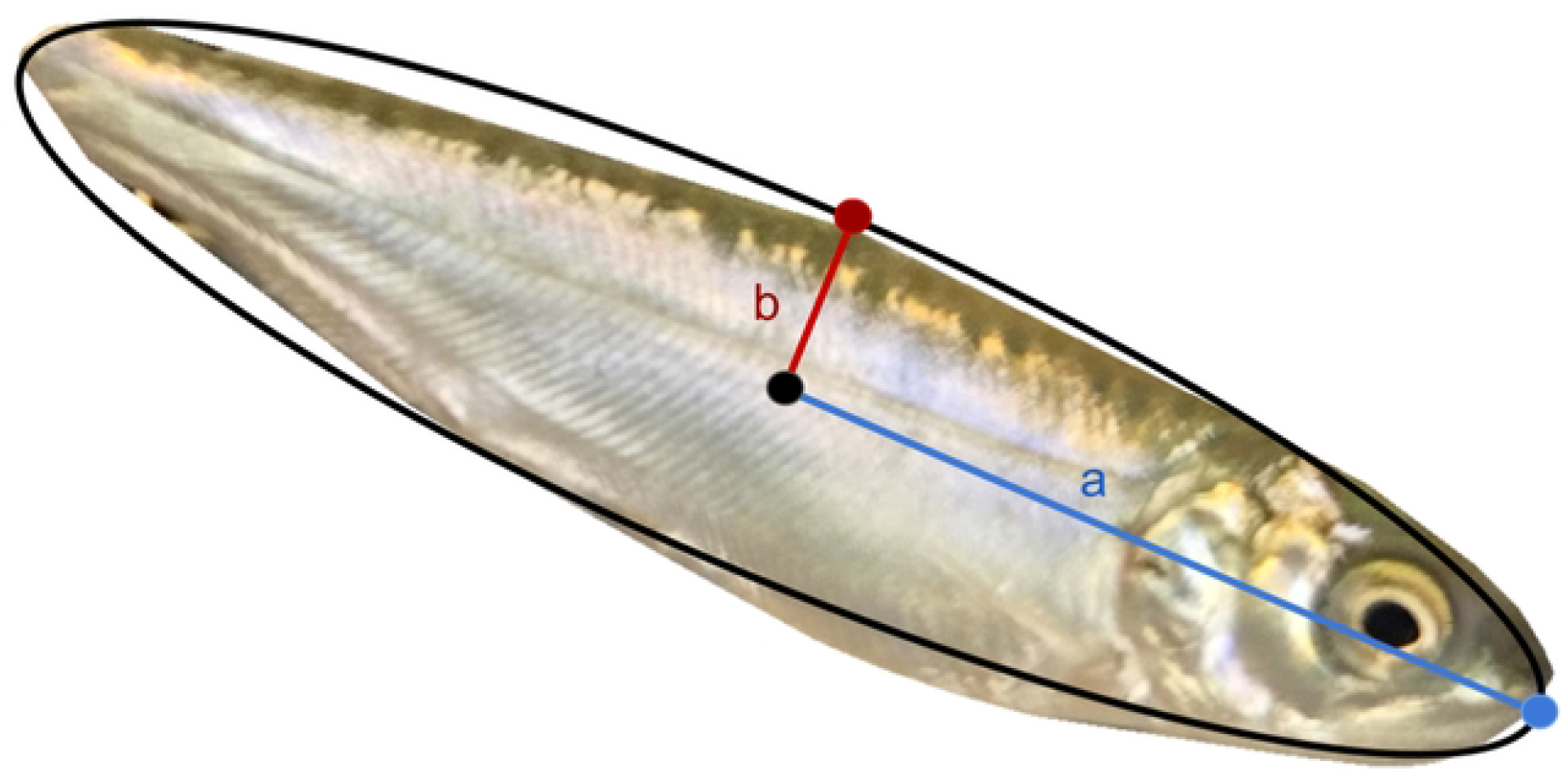
Ellipse of best fit for a given fish segmentation, with principal semi-axes labeled.

If the contour is well fit by an ellipse then the area of this ellipse, given by *A* = *πab*, should be well approximate the surface area measured. Thus this process could also be used to automatically check the success of the segmentation.

### 0.5 Weight Prediction Procedure

The dataset consisted of 149 images and ground truth weight measurements. A curated version of this dataset was also used, consisting of 109 images where low quality image or images with unwanted segmentation artifacts were removed. The HandsFreeFishing program was employed to measure many different morphometric features. Then, based on these features and additional modeling assumptions, several linear regression and multiple linear regression (MLR) models to predict juvenile fish weight. The MLR models were all trained in Python using the Scikit-Learn package.

## Results

For several weight prediction models, the shape of the fish body was assumed to be a 3D ellipsoid with semi-axes *a, b*, and *c*, denoting the horizontal semi-axis (length), the vertical semi-axis (height), and the depth semi-axis (girth), respectively, as depicted in Figure 7.

**Fig 7.**
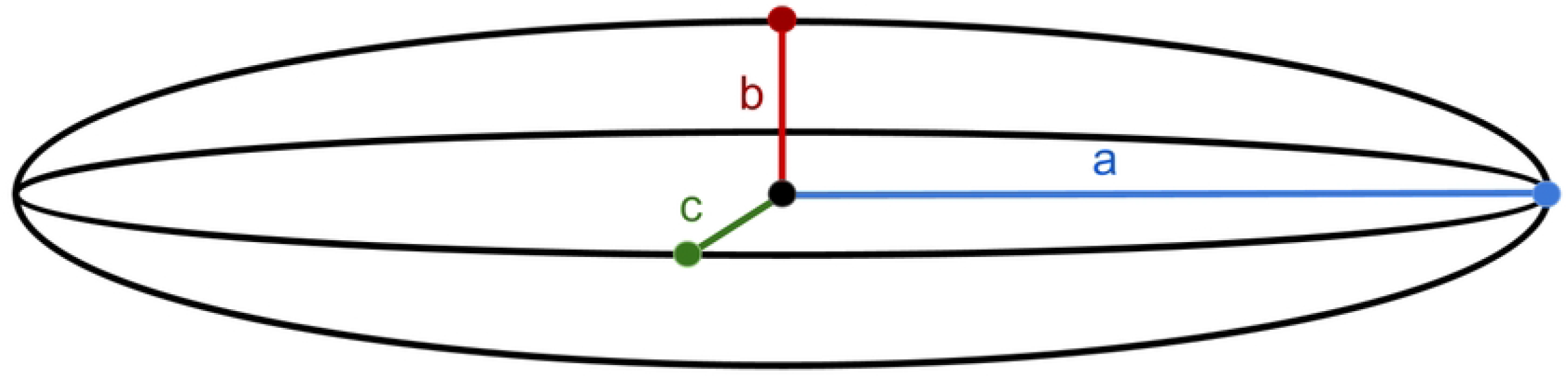
Ellipse of best fit for a given fish segmentation, with principal semi-axes labeled.

The volume *V* of an ellipsoid with these principal semi-axes is given by 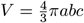, and that the area of a 2D ellipse with principal semi-axes *a* and *b* is given by *A* = *πab*. The surface area measured here is similar to the area of an ellipse, since the image is a 2D projection of the fish. Making the simplifying assumption that girth is directly proportional to height, it follows that *V* ∝ *SA* · *b*, and since volume should be directly proportional to weight, the quantities *V*_1_ = *SA* · *b* and *V*_2_ = *ab*^2^ should be directly related to weight via density. Further still, if it is assumed that height is directly proportional to length, i.e. all fish have the same proportions, then the model simplifies to *V* ∝ *b*^3^, and therefore 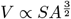. The quantities *V*_1_ and *V*_2_ and 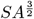 were all used in a standard 1D linear regression with the ground truth weight measurements to traine Model 1, Model 4, and Model 2, respectively, of which *V*_1_ and 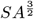 gave the most accurate results. The model believed to perform the best is Model 1, which achieved an *r*-squared statistic of 0.99, a mean average error of 0.16g, a mean average percentage error of 12% and an AIC score of-19.98. This is chosen over Model 2 even though Model 2 achieves a better AIC score on the cleaned dataset, because Model 2 implicitly assumes the length of the fish is proportional to the height and girth, which may be a bad assumption when there is more variance in the fish condition factors, for instance.

**Fig 8.**
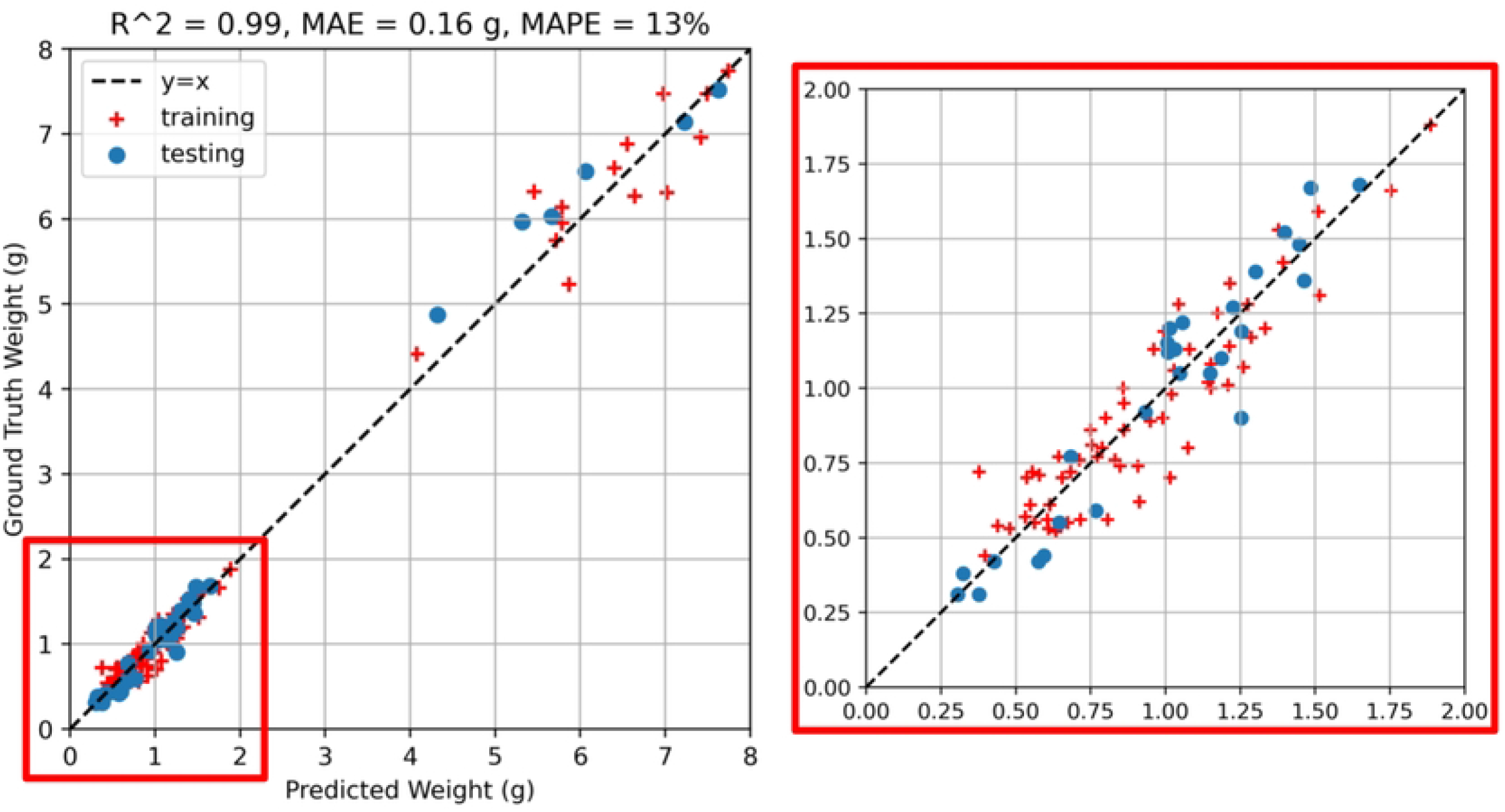
Multilinear regression between *SA, SA*^2^, and ground truth weight measurements. Model statistics shown are averages from 5 fold cross validation.

Several other models were considered as well. Model 3 is based on surface area, *SA*, and major and minor axis terms *a* and *b* repspectively. It uses these along with all second order combinations of *SA, a*, and *b* as explanatory variables, which consequently includes *V*_1_. This model achieved results similar to using *V*_1_ alone, but scored a worse in AIC and overall accuracy (Tables 1, 2). Model 5 uses the partitioned surface areas multiplied by the height estimate *b* as the explanatory variables. This model also performed well, but worse than simpler models in all categories. Finally, landmark length models, Model 6 and Model 7, are interesting to consider. They are trained using all the truss lengths computed as shown in Figure 5(c). Interestingly, considering these lengths cubed, which is motivated simply by volume being proportional to length cubed, the landmark lengths become much better predictors of weight. These models are summarized below:

- **Model 1**: a linear regression is performed with the quantity *V*_1_ = *SA* × *b*; this model implicitly assumes *b* ∝ *c*.
- **Model 2**: a linear regression is performed with the quantity 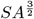; this model implicitly assumes *a* ∝ *b* ∝ *c*.
- Model 3: a multilinear regression is performed with the quantities *SA, a, b, SA*^2^, *SA* × *a, SA* × *b, a*^2^, *a* × *b, b*^2^; this model implicitly assumes *a* ∝ *b* ∝ *c*.
- Model 4: a linear regression is performed with the quanity *V*_2_ = *a* × *b*^2^; this model implicitly assumes *b* ∝ *c*, and further that *a* × *b* approximates the *SA* of the side profile of the fish well.
- Model 5: a multilinear regression is performed on the quantities *v*_*j*_ × *b*, where *v*_*j*_ is the surface area of the *j*th partition.
- Model 6: a multilinear regression is performed with each of the landmark lengths cubed;
- Model 7: a multilinear regression is performed with each of the landmark lengths;

**Table 1.**
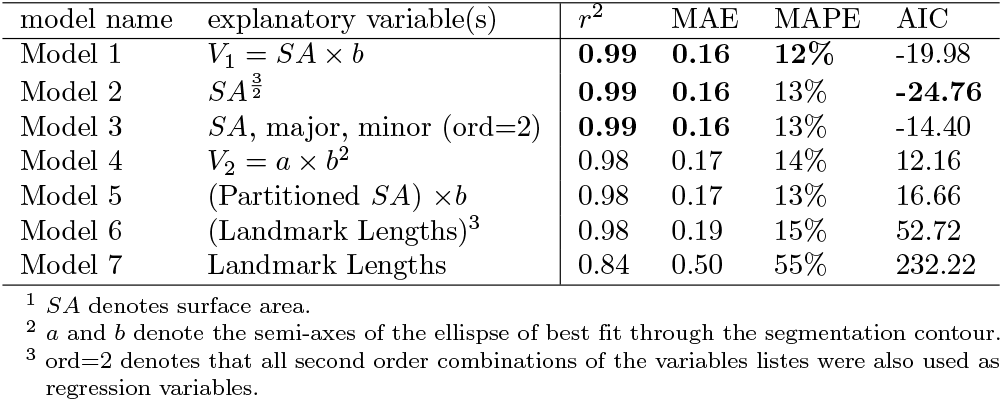
Cleaned dataset, 109 images.

| model name | explanatory variable(s) | $r^2$ | MAE | MAPE | AIC |
| --- | --- | --- | --- | --- | --- |
| Model 1 | $V_1 = SA \times b$ | **0.99** | **0.16** | **12%** | -19.98 |
| Model 2 | $SA^{\frac{3}{2}}$ | **0.99** | **0.16** | 13% | **-24.76** |
| Model 3 | $SA$, major, minor (ord=2) | **0.99** | **0.16** | 13% | -14.40 |
| Model 4 | $V_2 = a \times b^2$ | 0.98 | 0.17 | 14% | 12.16 |
| Model 5 | (Partitioned $SA$) $\times b$ | 0.98 | 0.17 | 13% | 16.66 |
| Model 6 | (Landmark Lengths)<sup>3</sup> | 0.98 | 0.19 | 15% | 52.72 |
| Model 7 | Landmark Lengths | 0.84 | 0.50 | 55% | 232.22 |

<sup>1</sup> $SA$ denotes surface area. <sup>2</sup> $a$ and $b$ denote the semi-axes of the ellipse of best fit through the segmentation contour. <sup>3</sup> ord=2 denotes that all second order combinations of the variables lists were also used as regression variables.

**Table 2.** Full dataset, 149 images.

| model name | explanatory variable(s) | $r^2$ | MAE | MAPE | AIC |
| --- | --- | --- | --- | --- | --- |
| Model 1 | $V_1 = SA \times b$ | **0.98** | **16%** | **0.13** | **9.69** |
| Model 2 | $SA^{\frac{3}{2}}$ | **0.98** | 0.17 | 14% | 13.79 |
| Model 3 | $SA$, major, minor (ord=2) | **0.98** | **0.16** | **0.13** | 26.14 |
| Model 4 | $V_2 = a \times b^2$ | 0.98 | 0.18 | 14% | 37.51 |
| Model 5 | (Partitioned $SA$) $\times b$ | 0.98 | 0.18 | 14% | 40.35 |
| Model 6 | (Landmark Lengths)<sup>3</sup> | 0.94 | 0.21 | 19% | 79190 |
| Model 7 | Landmark Lengths | 0.88 | 0.46 | 0.51 | 293.29 |

<sup>1</sup> $SA$ denotes surface area. <sup>2</sup> $a$ and $b$ denote the semi-axes of the ellipse of best fit through the segmentation contour. <sup>3</sup> ord=2 denotes that all second order combinations of the variables lists were also used as regression variables.

All weight prediction results are summarized in Tables 1 and 2.

The *r*-squared value, mean average error (MAE), mean average percentage error (MAPE), and Akaike information criterion (AIC) score is reported for each. Tables 1 shows results for a cleaned data set, where segmentations were manually inspected and images yielding poor segmentation results were discarded (109 images). As the goal of this model is to provide accurate weight data for unmeasured fish, this was done to achieve higher performance. On the other hand, Tables 2 second contains all 149 images, which is included to indicate that even with imperfect segmentation, accurate weight prediction is still feasible. All regression results are computed via 5-fold cross validation, using Scikit-Learn’s regression model and cross validation scoring function.

## Discussion

The HandsFreeFishing program extracted many important morphometric features from raw images of juvenile Chinook Salmon, including fork length, surface area, and eye diameter. This allows for the collection of important physical characteristics of juvenile salmon without the necessity of removing the fish from water, and the data extracted can be used for a variety of applications in fisheries ecology and management. Further, this work removes the current bottleneck in the morphometric data processing pipeline which will help expedite post processing of data and provide valuable morphometric datasets, including potential morphometrics that have not previously been measured.

Additionally, it addresses a growing concern with fish handling, as it should eventually allow a single in-water photo of an individual to be sufficient for measuring all important morphometric data. Fish handling often results in increased stress for fish [4]. Programs like this can help to minimize fish stress while still collecting critical metrics used to make management decisions. In addition to reduced handling and stress, additional metrics such as weight and condition factor can now be collected simultaneously in a single image. While working with threatened and endangered species these metrics can provide critical information to quantify success of various management actions while minimizing handling and stress on the fish.

For the purpose of predicting the weight of juvenile salmon, the program measured relevant features like surface area, fish height, and fork length. The actual weight of juvenile salmon were measured using a digital scale, with the juveniles ranging from 0.3 to 7.8 grams. Several linear regression models were then trained to predict weight from the measured features. The most successful models assumed that a fish body can be well approximated by an ellipsoid in 3D, who’s girth is proportional to its height. From this assumption it follows that the volume of the fish body is proportional to the surface area multiplied by the height. Therefore, by predicting both surface area and the fish height, the weight can be estimated accurately. This is verified empirically here, as Model 1 obtained the best prediction results and second best AIC score.

Collecting more ground truth weight data for the predictive models to be trained on will allow for more accurate and potentially more complex models. Model 5 and Model 6, for example, suffer due to the size of the training dataset, and could potentially outperform Model 1 if trained on a larger number of fish. Other future work with this program could involve new feature extraction methods and algorithms for accurate anatomical landmark point placement. For example, no attempt has yet been made to segment the midline, which could help with orienting the fish and improve the fork length estimate. Predictive models of other morphometric measurements, such as condition factor, would be an interesting direction to pursue.

## Conclusion

By adapting a state of the art segmentation model like Meta’s Segment Anything Model [6], highly accurate segmentations of juvenile Chinook Salmon are readily attained. Relying on this accurate segmentation data, the HandsFreeFishing program is able to extract a wide variety of morphometric features automatically. The approach taken here, and a lot of the software and methods developed, should be adaptable to other species and life stages, and the use of the HandsFreeFishing program to create a growing database of these morphometric features is an exciting prospect and highly encouraged.

## Supporting information

### S1 Appendix. Fin and Eye Segmentation Heuristics

Here the heuristics used to segment individual fins and the eyball of the fish are discussed. Once the initial fish contour is extracted and rotated, the left most point of the contour lies close to the snout of the fish, and the right most point lies somewhere along the caudal fin. Let *c* = [*c*_0_, *c*_1_, … *c*_*n*_] where *c*_*i*_ = (*x*_*i*_, *y*_*i*_) for 0 ≤ *i* ≤ *n* represent this rotated contour. The fork length is then computed as outlined in 0.4. For the following heuristics, the quantity *x*_*max*_ = max_0≤*j*≤*n*_ *x*_*j*_ will be used frequently. This represents the largest *x* value and thus the right most point of the contour. The average *x* value 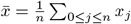 will also be utilized. Note that these countour points are extracted from the fish segmentation, and thus small *y* values are higher in the image, whereas large *y*-values are lower in the image. Small *x* values correspond to the left side of the image whereas large *x* values correspond to the right side of the image.

1. (*Dorsal Fin*) To estimate a bounding box for the dorsal fin, all points of the contour *c* for which 0.5*x*_*max*_ ≤ *x*_*i*_ ≤ 0.6*x*_*max*_ are found. From these points, the largest *y*-value is computed, and this is taken to approximate the top most point of the dorsal fin. After this point is located, a bounding box with a width of 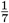 the fork length and height of 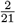 the fork length is centered about this point and given to the segmentation model, which produced the dorsal fin segmentation.
2. (*Adipose Fin*) To estimate a bounding box for the dorsal fin, all points of the contour *c* for which 0.7*x*_*max*_ ≤ *x*_*i*_ ≤ 0.8*x*_*max*_ are found. From these points, the smallest *y*-value is computed, and this is taken to approximate the top most point of the adipose fin. After this point is located, a square bounding box with a width of 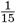 the fork length and height of 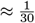 the fork length is centered about this point and given to the segmentation model, which produced the adipose fin segmentation.
3. (*Caudal Fin*) To estimate a bounding box for the caudal fin, first all points of the contour *c* = [*c*_0_, *c*_1_, … *c*_*n*_] for which 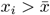 were computed. These are the contour points along the back half of the fish where the caudal fin is. Next, the minimum and maximum *y*-values along this subset of the contour were computed. A ratio of 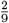 is provided, which estimated that caudal fin will contained in the last 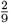 of the fork length. Finally, a bounding box which is vertically between the minimum and maximum *y* values, and horizontally between the right most point of the contour and the last 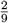 of the contour was given to the segmentation model, which produced the caudal fin segmentation.
4. (*Anal Fin*) To estimate a bounding box for the dorsal fin, all points of the contour *c* for which 0.6*x*_*max*_ ≤ *x*_*i*_ ≤ 0.7*x*_*max*_ are found. From these points, the largest *y*-value is computed, and this is taken to approximate the bottom most point of the anal fin. After this point is located, a square bounding box with a width and height of 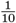 the fork length is centered about this point and given to the segmentation model, which produced the adipose fin segmentation.
5. (*Pelvic Fin*) To estimate a bounding box for the dorsal fin, all points of the contour *c* for which 0.5*x*_*max*_ ≤ *x*_*i*_ ≤ 0.6*x*_*max*_ are found. From these points, the largest *y*-value is computed, and this is taken to approximate the bottom most point of the pelvic fin. After this point is located, a bounding box with a width of 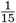 the fork length and height of 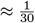 the fork length is centered about this point and given to the segmentation model, which produced the pelvic fin segmentation.
6. (*Pectoral Fin*) To estimate a bounding box for the caudal fin, first all points of the contour *c* = [*c*_0_, *c*_1_, … *c*_*n*_] for which 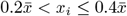 were computed. From these points, the maximum *y* value was computed. After this point is located, a square bounding box with a width and height of 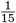 the fork length is centered about this point and given to the segmentation model, which produced the pectoral fin segmentation.
7. (*Eyeball*) To estimate a bounding box for the caudal fin, first all points of the contour *c* = [*c*_0_, *c*_1_, … *c*_*n*_] for which 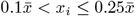 were computed. From these points, the mean *y* value was computed to estimate the midpoint which would be around eye level. After this point is located, a square bounding box with a width and height of 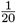 the fork length is centered about this point and given to the segmentation model, which produced the pectoral fin segmentation.

These heuristics are included for the sake of completeness, but also to emphasize that these procedures are easily altered and customizable to the particular interests of the researcher. For the purposes of this manuscript, the main goal was to disclude fins whenever present, and accurate fin segmentation was not of critical importance. Hence these heuristics are not optimized, and could be improved if the main task was to estimate caudal fin sizes, for example.

## Acknowledgments

We would like to thank Chief Sisk, Marine Sisk, and the Cultural Resource Specialists of the Winnemem-Wintu Tribe for pushing us to innovative in a collaborative space. The California Department of Fish and Wildlife funded the development of this project (agreement Q2396019). Dr. Jay Lund for supporting development. We are grateful to Matt Salvador for data collection and all who provided feedback on the development of this program.

---

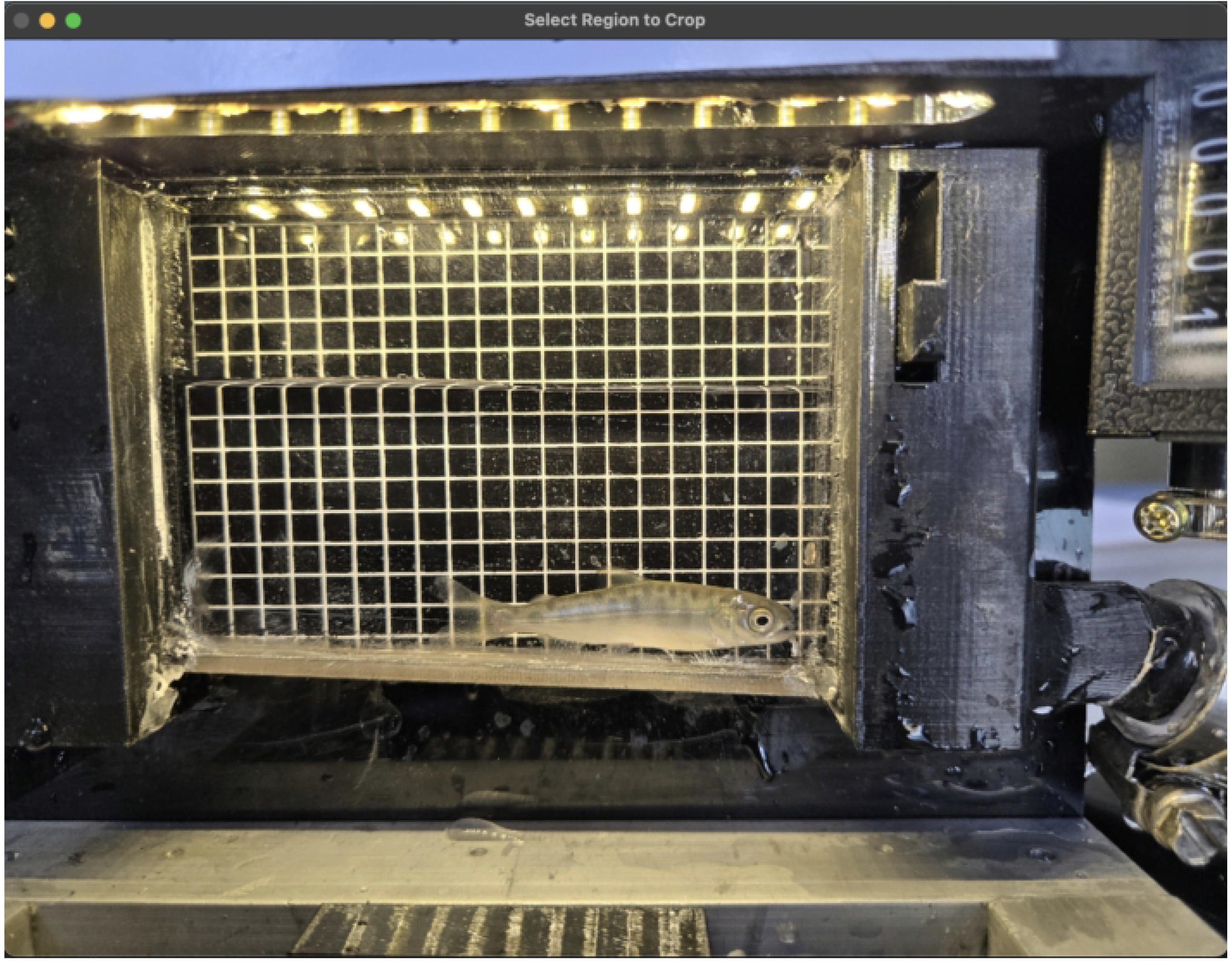

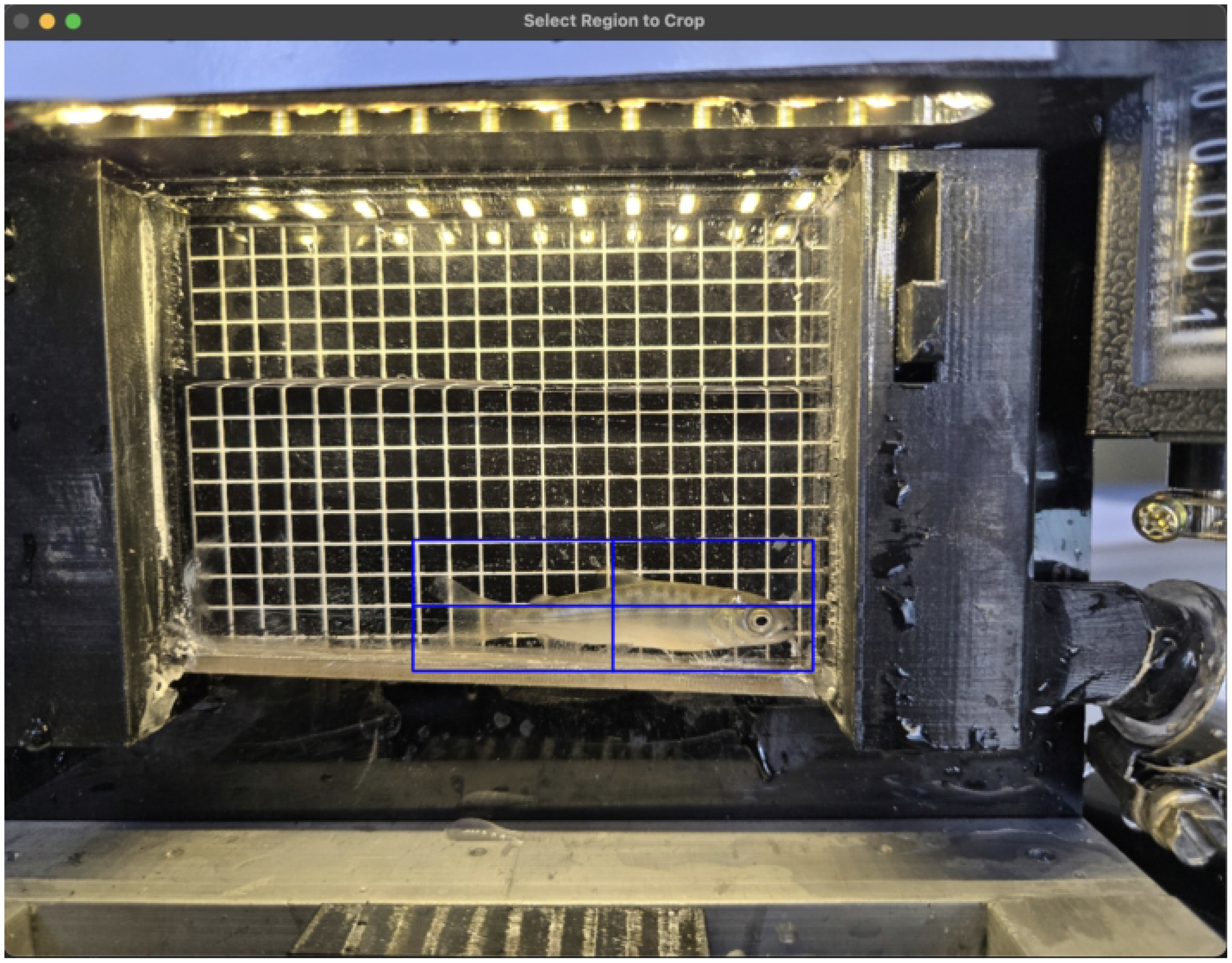

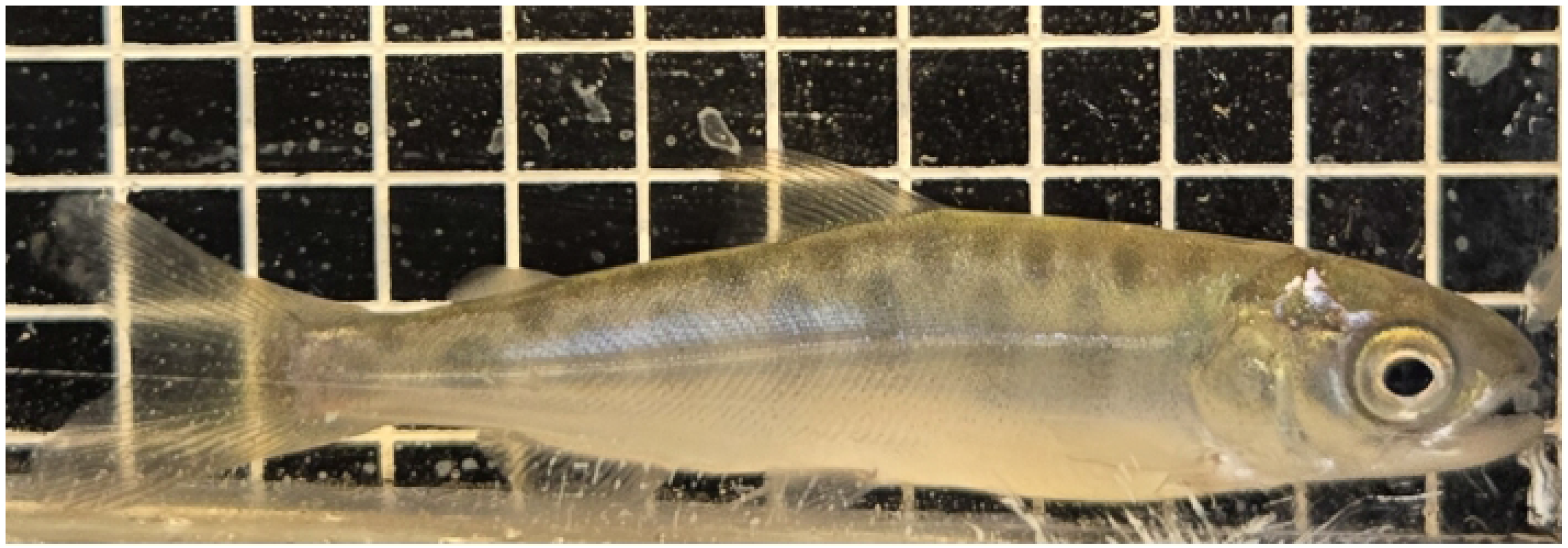

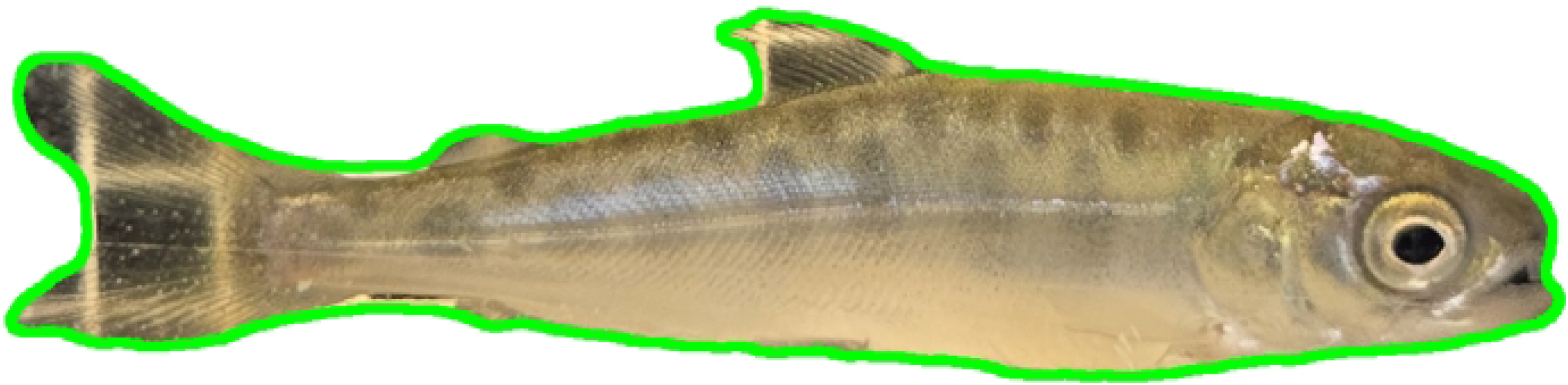

## Notes

### Competing Interest Statement

The authors have declared no competing interest.

